# Drivers of systemic male-female allele frequency divergence in humans

**DOI:** 10.64898/2026.01.06.698007

**Authors:** Matthew Ming, Arbel Harpak

## Abstract

Allele frequency differences between males and females in genetic studies have been interpreted as suggestive of sex differences in natural selection. However, systemic differences may also arise through sex differences in study participation as well as bioinformatic artifacts. To mitigate these confounding effects, we performed a meta-analysis of sex differences in allele frequencies across three genetic studies. We identify twelve concordant genes with cross-study evidence of sex differences in allele frequency. Using previous literature about the twelve candidates, we propose four hypotheses for how and when sex differences in allele frequencies arise and suggest which hypotheses are plausible for each candidate. For example, beyond candidates that plausibly reflect sex differences in viability, we also find candidate loci potentially underlying fitness differences between X/Y-carrying sperm. Taken together, our results offer a clearer delineation of the factors driving sex differences in allele frequencies, including the timing and mechanisms of natural selection.

## Introduction

Sex-differential selection (SDS), where selection strengths differ between males and females^1,2,3^, could drive sex differences in allele frequencies. However, searching for between-sex allele frequency divergence is underpowered, even with modern biobank sample sizes, because the independent assortment of chromosomes during gametogenesis homogenizes allele frequencies. The association of autosomal alleles with the X (Y) chromosome or XX (XY) karyotype resets in each generation. Therefore, researchers have searched for signals of SDS by leveraging information about alleles subject to selection and by integrating signals of allele frequency divergence from variants across the genome that are hypothesized to reflect the selection process in a single generation^1,4,5^.

To our knowledge, all previous studies investigating SDS generally did so using data from a single genetic study^2,5,6,7^. However, using a single genetic study for large-scale genomic analysis limits interpretation of the results because of study-specific biases and artifacts. Factors that drive study participation can differ by sex^8,9^ without being related to viability. Although other studies have found no sex-specific association of traits with UK Biobank (UKB) participation^10^, the possibility of sex-specific participation biases affecting observed between-sex allele frequency differences^9^ remains a major concern. In addition, bioinformatic artifacts^11,12^ can drive misalignment or mis-mapping of autosomal alleles to sex chromosomes, or sex-chromosomal alleles to autosomes^4^, and may in some cases be study-specific due to different sequencing technologies, variant calling procedures, and other technical factors.

Here, we use a meta-analysis approach to mitigate study-specific confounders that limit the interpretation of sex differences in allele frequency as indicative of SDS. We analyze allele frequency differences in the Genome Aggregation Database (gnomAD)^13^, UKB^14^, and All of Us (AoU)^15^. We identify significant signals that replicate across studies, which we therefore view as offering more robust evidence of SDS than previous work. Specifically, we find twelve candidate genes putatively under SDS. Finally, we delineate hypotheses for possible selective mechanisms through which this differentiation is driven.

## Results and Discussion

### Identifying concordant male-female allele frequency differences across studies

Under contemporary SDS, we expect allele frequencies to diverge between males and females. We tested whether sample allele counts are consistent with a null hypothesis of random distribution of alleles between males and females due solely to drift and randomness in study participation. Using a Chi-squared test with 1 degree of freedom we tested across autosomal biallelic single nucleotide polymorphisms (SNPs) in two non-overlapping datasets: the UK Biobank (UKB) and the Genome Aggregation Database (gnomAD). We only considered SNPs within gene bodies or up to 1kb upstream of a transcription start site as annotated in Ensembl^16^. We obtained gene-level p-values by considering the lowest p-value among all SNPs associated with each gene (henceforth, we term the corresponding SNP the “lead SNP” for each gene-study pair). We focused on the top 1% of the most significant genes of each study sample (**Fig. S1**). Out of 19,881 protein-coding genes, we identified 343 genes that were significant in either UKB or gnomAD, with 21 genes significant in both datasets.

For these 21 overlapping significant genes, we calculated the evidence for allele frequency divergence (again using a chi-squared test) in a third dataset, AoU. We also constructed a matched null distribution of p-values in AoU using 200 randomly selected genes across the genome (excluding the 21 candidates). We additionally considered genes as significantly suggestive of SDS in AoU if their p-value was lower than the 1^st^ percentile of this empirical null.

In total, we identified twelve genes with concordant evidence for sex differences in allele frequencies across all three datasets (**Table 1**). We refer to these genes as “cross-study significant genes”. We note that the lead SNP for almost all of the cross-study significant genes was located within an intron. The one exception was the lead SNP for DPEP2 in UKB (chr16:67994261), which was a non-synonymous mutation of Glutamine to Glutamic Acid at codon 127 (Q127E).

**Table 1:** Hypothesis concordance for significant genes across three studies. We identify genes with evidence for male-female allele frequency differences across three studies. DPEP2 is the only gene with a non-synonymous lead SNP in one study (UKB); all other lead SNPs were intronic. Significant deviation from Hardy-Weinberg Equilibrium within a sex is consistent with selection acting in that sex. If a lead SNP tags the same selected variant across studies, we expect concordant signs of male-female allele frequency difference across studies. Here, we focus on the sign concordance at the SNP in highest summed pairwise LD with the three study-specific lead SNPs. Asterisks indicate genes that we view as concordant with multiple hypotheses. The highlighted gene ANKRD36C represents the only gene with cross-study significance replicating in a sensitivity analysis (*MAGMA* analysis).

| Associated Hypothesis | Gene | Deviation from HWE Males | Deviation from HWE Females | Sign Concordance (Highest LD SNP) | Functional Notes |
| --- | --- | --- | --- | --- | --- |
| Spermatogenesis and fertilization | ANKRD36 | AoU only | UKB and AoU | 3/3 | Overexpressed in Testes<br>Associated with a network of proteins important in sperm maturation, motility, and capacitation (Kant et al., 2019)<br>Protein product found in seminal fluid and likely to play role in male fertility (Milardi et al., 2012) |
|  | ANKRD36C | UKB and AoU | UKB and AoU | 2/3<br>UKB discordant | Overexpressed in Testes<br>Protein product found in seminal fluid and likely to play role in male fertility (Milardi et al., 2012) |
|  | GRID2* | UKB and AoU | Not significant | 2/3<br>UKB discordant | Overexpressed in Testes, especially elongated spermatids and early spermatids (Human Protein Atlas)<br>GWAS shows association with Azoospermia and Oligospermia (Chatziparasidou et al., 2024) |
|  | DUS2 | Not significant | Not significant | 3/3 | Specific 3kb transcript only found expressed in testes (Kato et al., 2005) |
| Fetal development and Viability | CNTNAP2* | UKB and AoU | UKB and AoU | 3/3 | Involved in neuronal cell adhesion<br>Associated with many neurological disorders including Autism Spectrum Disorder, Schizophrenia, Intellectual disability, Dyslexia, and Language impairment (Rodenäs-Cuadrado, Ho, and Vernes, 2013)<br>Associated with sexually dimorphic social disruption in KO mice (Dawson et al., 2023)<br>Genetic variants associated with gender differences in dyslexic children (Gu et al., 2018)<br>Sex-specific effects of CNTNAP2 mutations in visual cortical brain area (Townsend and Smith, 2017) |
|  | LAMA2 | Not significant | Not significant | 3/3 | Associated with congenital muscular dystrophy, but this form of the disease is not known to have significant sex differences (Tan et al., 2021) |
|  | SVEP1* | AoU Only | AoU Only | 2/3<br>gnomAD discordant | Associated with lymphatic and cardiovascular vessel development in mouse and zebrafish models (Marooka et al., Circ Res, 2017; Kerpanen et al., Circ Res, 2017)<br>The exact mechanism through which SVEP1 acts during development are still not well understood (Elenbaas et al., Trends Mol Med, 2023) |
| Post-natal disease and viability | RUNX1 | Not significant | Not significant | 3/3 | Associated with acute myeloid leukemia (AML), where RUNX1 mutations were found to be more common in males with AML, and mutations led to worse remission outcomes in females (Ozga et al., 2023) |
|  | GRID2* | UKB and AoU | Not significant | 2/3<br>UKB discordant | Associated with spinocerebellar ataxia type 18 (Panda, Sharawat, and Dawman, 2020), a very rare form of ataxia where individuals suffer progressive incoordination of gait, hands, and eyes (Rossi et al., 2014) |
|  | PTPRM | UKB and AoU | UKB and AoU | 3/3 | High expression associated with poor prognosis in cervical cancer (Liu et al., 2020)<br>Methylation patterns associated with colorectal cancer (Laczanska et al., 2013) |
|  | DCC | UKB only | Not significant | 3/3 | Deleted in Colorectal Cancer, associated with colorectal cancers (Fearon et al., 2020; Keino-Masu et al., 1996) |
|  | DLG2 | Not significant | AoU only | 2/3<br>UKB discordant | Deletion associated with neurodevelopmental disorders (Chen et al., 2024)<br>Tumor Suppressor in osteosarcoma (Shao et al., 2018)<br>Deficiency leads to reduced sociability and increased repetitive behavior in mice (Yoo et al., 2020)<br>Associated with pubertal disorders (Jee et al., 2020) |
|  | DPEP2 | Not significant | Not significant | 2/3<br>AoU discordant | Associated with outcomes in lung adenocarcinoma (Han et al., Sci Rep 2023; Wang et al., 2023)<br>Associated with risk of coronary artery disease and T2 diabetes (Giufrida et al., 2024) |
|  | SVEP1* | AoU Only | AoU Only | 2/3<br>gnomAD discordant | Associated with several diseases and lifetime health by numerous studies, including Coronary Artery Disease, Hypertension, Type 2 Diabetes, Dementia, and Glaucoma (Elenbaas et al., Trends Mol Med, 2023) |
| Neurobehavior and participation | CNTNAP2* | UKB and AoU | UKB and AoU | 3/3 | Involved in neuronal cell adhesion<br>Associated with many neurological disorders including Autism Spectrum Disorder, Schizophrenia, Intellectual disability, Dyslexia, and Language impairment (Rodenäs-Cuadrado, Ho, and Vernes, 2013)<br>Associated with sexually dimorphic social disruption in KO mice (Dawson et al., 2023)<br>Genetic variants associated with gender differences in dyslexic children (Gu et al., 2018)<br>Sex-specific effects of CNTNAP2 mutations in visual cortical brain area (Townsend and Smith, 2017) |

\* We view multiple hypotheses as equally plausible for this gene

We recognized that increased gene length may correlate with decreased p-value for the lead SNP in the gene. In addition, non-coding regions may affect genes beyond the nearest one, including interactions between introns in one gene and other genes in *trans*^17^. Therefore, large genes—which by extension will have larger intronic regions—could show inflated p-values. Indeed, we observed that gene-level p-value tends to increase with gene size, and most of the cross- study significant genes we identified are among the top 10% longest genes in the genome (**Fig. S4**). Therefore, we additionally used *MAGMA* to summarize signals across SNPs.

*MAGMA* aggregates SNP-level statistics (in our case, SNP-level Chi-squared test statistics) from SNPs within a gene and summarizes them into a gene-level test statistic, accounting for linkage disequilibrium among gene-associated SNPs (**Methods**) and gene size. Using this method, only one gene—ANKRD36C—remained cross-study significant. ANKRD36C was significant in all three studies, while ANKRD36 was significant in UKB and AoU but not gnomAD. Two other genes, GRID2 and DCC, were significant in AoU only (**Table S1**). Visualizing the between-sex allele frequency divergence for these genes, some qualitative differences between the *MAGMA*- significant and the nine other cross-study significant genes became more apparent. In ANKRD36 and ANKRD36C, many sites in close proximity have strong signals of concordant divergence, a “hitchhiking” response one would expect even when selection only impacts differentiation in a single generation^2,19,20,21^ (**Fig. 1, S2a**). In other genes, one or two nearby SNPs show significant allele frequency differentiation (**Fig. S2**) with otherwise little evidence of hitchhiking. Since ANKRD36 and ANKRD36C are paralogs, we were concerned with spurious signals resulting from interlocus gene conversion^22,23^ or bioinformatically, from mismapping. While we could not rule these options out, we found that the paralogous 22kb regions surrounding the lead SNPs in each gene are somewhat diverged (identity = 82%), reducing the concern of either interlocus gene conversion^23,24^ or mismapping.

**Figure 1:**
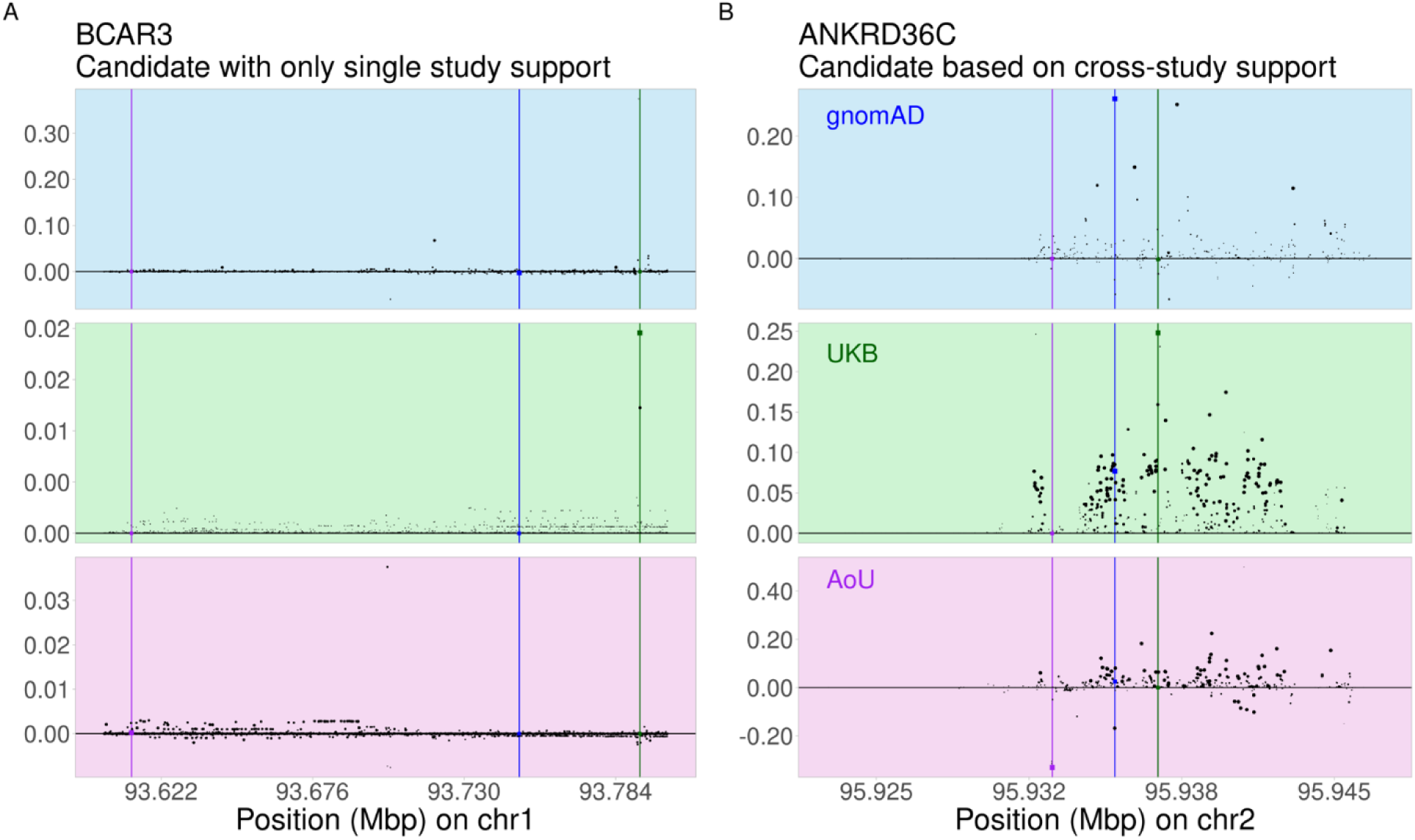
Concordant, large allele frequency differences across biobanks indicate genes under sex-differential selection. We identified genes with concordant significant between-sex allele frequency differentiation across datasets. Here, we depict the most significant SNPs in BCAR3 (A, an example of a non-cross-study significant gene) and ANKRD36C (B, a cross-study significant gene), as well as 10kb up- and downstream of the top SNPs. Each point is a single SNP, with its size reflecting Chi-squared test p-value. The region plotted falls within the respective gene body. The y-axis shows male-minus-female allele frequency, polarized by the allele with higher frequency in males in UKB. Background colors correspond to the dataset analyzed. Colored square points represent the lowest p-value site in each dataset, with a line of matching color showing the position of these points.

Given the lack of an archetypic hitchhiking pattern (expected at selected sites^2,19,25^) around lead SNPs at most candidate genes, we considered the possibility of a bioinformatic artifact that impacts all studies. However, we note that the genotyping technology and bioinformatic pipelines vary among these studies. In addition, artifacts such as mis-mapping to/from sex chromosomes^4^ should result in a hitchhiking pattern as well. Therefore, while the evidence is most compelling for ANKRD36C, we still consider the other eleven genes as potential targets of SDS moving forward.

If a SNP is under SDS, we expect the same allele to be more frequent in males than females across datasets. We examined the sign concordance—whether the same allele is more common among males than among females—across studies, focusing on the lead SNPs. We initially performed six pairwise comparisons per cross-study significant gene, where the sign of the lead SNP in one of the three studies was compared with the sign in the other two (even if it was not the most significant SNP in the other two datasets). The sign was often inconsistent when focusing on the lead SNPs individually, but this inconsistency appeared to be noise-driven rather than indicative of an overall strong sign concordance in cross-study significant genes (**Figs. 1, S2**). We therefore used an alternative approach to capture composite sign concordance. Specifically, we examined the sign concordance in the SNP with the strongest combined pairwise LD with all three lead SNPs (“Linked site cross-study sign concordance,” **Methods**). Using this method, we found ANKRD36C showed discordant sign in UKB. We observed sign concordance for almost all other non-MAGMA cross-study significant genes, with seven of the eleven showing concordant signs for all studies (**Table 1**).

SDS may occur through one of several distinct modes. For example, selection may be sex-antagonistic (different alleles being favored in males vs. females), sex-specific (selection is only acting on one sex), or sex-concordant but with different strengths of selection between sexes (the same allele is favored in both sexes but more strongly beneficial in one). To classify the support for different modes of SDS, we evaluated the sex-specific deviation from Hardy-Weinberg Equilibrium (HWE) sample frequencies using a Fisher’s Exact Test, with the rationale that deviations within a sex are consistent with selection acting on that sex^7,23^, and that the test is independent of the distribution of genotypes in the other sex. We therefore tested in males and females separately and in AoU and UKB separately. (We could not test in gnomAD since the test required genotype counts which were not publicly available). Several of the genes showed significant deviations from HWE that replicated across studies ( **Fig. S5**): GRID2 and DCC in males only, and DLG2 in females only (consistent with sex-specific selection); and three genes— CNTNAP2, PTPRM, and ANKRD36C—in both sexes (consistent with either sex-antagonistic or sex-concordant selection). We note that the direction of deviation from HWE in all cases was towards underabundance of heterozygotes (**Fig. S5, Table 1**), consistent with directional selection within each sex.

### When in life does selection act differently in males and females?

SDS may operate in different phases between gametogenesis and sampling in adulthood ( **Fig. 2**). Specifically, we hypothesize that allele frequency divergence can arise as a result of 1) selection on gametes prior to and during fertilization, 2) viability selection during fetal development, 3) post-natal viability selection through adulthood, or 4) sex-biased differences in genetic study participation. In **Table 1**, we categorize the cross-study significant genes by hypotheses with which they seemed to align most closely with based on existing literature. We expand on this possible alignment below. However, it is important to note that these hypotheses are largely speculative, and the following section seeks only to illustrate what we believe to be plausible classification of drivers towards candidate-specific hypotheses.

**Figure 2:**
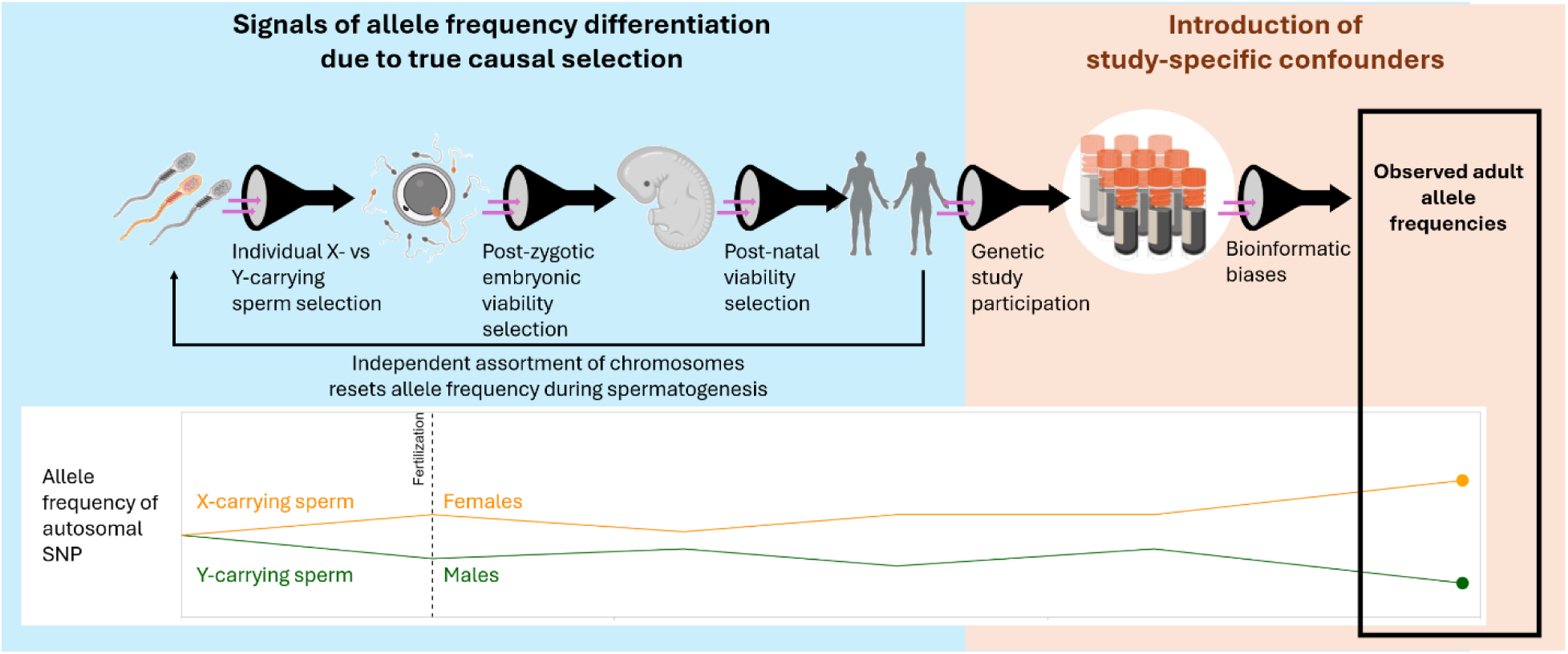
Hypotheses for drivers of between-sex allele frequency divergence. We consider five potential drivers of autosomal allele frequency divergence between males and females in a genetic study sample.

### Selection in Sperm

Some of the cross-study significant genes are highly expressed in male reproductive cells and organs (among other tissues). For example, GRID2, ANKRD36, ANKRD36C, PTPRM, and DCC are expressed in the testes and late spermatids^27,34^, and DUS2 has a splicing transcript form that is only expressed in testes^28^. While studies have shown that male infertility is often associated with increased mortality^29,30^, it is unclear if the cross-study significant genes we have identified are involved in this decrease in viability or in pathways of comorbidity. These genes should also be under weak or no selection in females, and the frequency in females should fluctuate due only to drift, so sex differences between studies should be inconsistent if viability selection is the primary driving force. This raises the question: how could genes which primarily affect male spermatogenesis contribute to sex-differential selection if they should experience no selection in females? One possible explanation is that selection can act on individual sperm carrying different sex chromosomes. For example, if an autosomal allele impacts sperm motility differentially in individual sperm carrying different sex chromosomes, this would lead to sex differences in allele frequencies among XX and XY zygotes due to differential fertilization rates.

For this hypothesis to hold merit, two conditions need to be fulfilled. First, sperm would need to have individual-level phenotypic variation that ultimately affects its likelihood of fertilization; and second, there would be a genetic component of sperm phenotype which is based on individual sperm haplotype rather than the genotype of the adult male producing the sperm. Regarding condition one, individual sperm generally are known to vary significantly by tail length, head size, motility, survivability, and many other phenotypes^31,32^. Some studies report differences between these physical characteristics in X- and Y-carrying sperm, while other studies dispute these findings, so systemic X- versus Y-sperm differences remain contentious^33^. However, physical characteristics are not the only characteristics that can increase likelihood of fertilization. Increased abundance of sperm carrying specific alleles would also increase the likelihood of fertilization by those sperm. If an autosomal allele causes increased sperm abundance only when paired with a specific sex chromosome, this will lead to differences in allele frequency in male and female adults. GRID2, one of the cross-study significant genes, encodes a glutamate receptor protein. Mutations in GRID2 are associated with low sperm count^34^. This decreased sperm count could be due to the effect of an existing allele on the meiotic or spermatogenic process, or due to a *de novo* mutation which arose during spermatogenesis. Recent work has shown that some mutations may drive increased replication in sperm precursors, leading to an enrichment for these mutations in sampled adults^35^. Regardless of the mechanism driving allele frequency differences in sperm, differences in sperm phenotype can lead to differences in individual sperm fertilization success.

Regarding condition two, the extent to which sperm genotype affects its phenotype is also contentious. During meiosis, sperm precursor cells form cytoplasmic bridges that allow gene products to be shared between neighboring sperm^36^. Although each sperm has its own haplotype, the mRNA or proteins available to each sperm may not be solely determined by that one sperm’s alleles but also by those of its neighbors. We found that six out of our cross-study significant genes were expressed in sperm, representing significant enrichment compared to all genes (enrichment ratio = 3.28, p = 0.0050); further, two out of four of our cross-study significant spermatogenesis-related genes (GRID2 and ANKRD36C) are not shared across these cytoplasmic bridges, based on comparison with a study of which proteins are shared across bridges^37^ (**Table S2**). This means the specific allele for these genes carried by an individual sperm should reflect the gene products in that sperm, and there should thus be a direct connection between sperm phenotype and haplotype. Selection acting on the sperm phenotype can then work to effectively drive between-sex differences in allele frequencies in the offspring.

This hypothesis operates under the assumption that X- and Y-carrying sperm are differentially selected prior to fertilization. However, the fertilizing sperm continues to have an effect on the zygote. Sperm deposit proteins into the neo-zygote at the moment of fertilization, and these deposits are known to have some effect on the zygote’s viability^38^. If these deposits act differently based on whether the sperm also contributed an X or Y chromosome, it could affect adult allele frequencies. Although this form of selection technically falls under post -zygotic viability selection (see the following section), we mention it here because the viability of the zygote at this stage is driven by sperm-specific transcripts.

### Post-zygotic viability selection

Autosomes segregate independently from the sex chromosomes. Therefore, differences in allele frequencies between males and females are expected to revert to zero each generation. Male- female allele frequency divergence above random sampling variance has therefore been assumed to primarily reflect differences in viability selection arising after conception^1,2,5,39,40^. To evaluate the potential support for this hypothesis among cross-study significant genes, we further stratify this hypothesis into two groupings according to the timing of viability selection: sex-differential lethality during fetal development and lethality due to disease post-birth.

Studies suggest somewhere between 40-60% of all human embryos die between fertilization and birth^41^ and around 1 in 200 newborns die annually in the United States, with 40% of deaths being due to congenital developmental diseases^42^. Genes associated with pre-natal development are expected to play a role in fetal and neonatal viability and may experience selection leading to allele frequency shifts in adults. An example of a top significant gene which affects fetal development is LAMA2. This gene encodes an extracellular protein which serves to organize cells during embryonic development. Mutations in LAMA2 are also associated with forms of muscular dystrophy including LAMA2-related congenital muscular dystrophy (LAMA2- CMD) and limb-girdle muscular dystrophy-23^43^, both of which have observable symptoms and negative effects in newborns. LAMA2-CMD most commonly presents as an early-onset form where symptoms are generally already manifested at time of birth or within the first few months of life^44^. Lung infections due to complications from breathing muscle weakness are often observed, with mortality from these symptoms having been observed in 50% of patients^43^. Despite strong signals for male-female allele frequency divergence in our analysis, there is no strong evidence for consistent male-female differences for LAMA2-CMD in humans^43,45^. However, there was evidence for significant sex differences in the effects and progression of LAMA2-CMD-related symptoms, as well as response to an experimental treatment, in mice^46^.

There are more known sex differences in post-natal disease. Several cross-study significant genes are associated with adult-onset diseases. Acute myeloid leukemia (AML) is a blood cancer that accounts for the highest percentage of leukemia deaths^47^. Unlike LAMA2-CMD discussed above, for which symptoms can be observed at birth, the age of onset and diagnosis of AML is in mid-to-late adulthood^48^. RUNX1, one of the cross-study significant genes, is involved in hematopoietic stem cell differentiation^49^ and is associated with development of AML^50^. Interestingly, the form of AML associated with RUNX1 mutations appears to differ in its effect between males and females. Males carrying RUNX1 mutations are more likely to have AML than RUNX1-mutated females^44,51^, and RUNX1 mutations were associated with worse remission outcomes in females but not in males^52^. Several other types of cancer are associated with mutations in the PTPRM gene. PTPRM codes for a protein that is a member of the tyrosine phosphatase family, a group of signaling molecules known to regulate a variety of important cellular functions^53^. High expression of PTPRM has been linked with poor prognosis in cervical cancer^54^, which only occurs in females. Hypermethylation of PTPRM is also associated with colorectal cancer^55^, a form of cancer more prevalent and with higher mortality in men^56^.

For diseases occurring during fetal development such as LAMA2-CMD, pre-birth death or mortality in newborns up through adolescence will lead to differences in a population when sampled at conception and in adulthood. Even for diseases occurring during adulthood, if death occurs before sampling, we would still see this selective pressure reflected in our observed allele frequencies. As several genes are identified that have known associations to disease, we suggest that viability selection from post-zygotic development and continuing through adulthood can lead to observed differences in allele frequencies by the time individuals are sampled.

Finally, we note a marginal enrichment of associations with esophageal cancer, leukemia and colorectal cancer among the 12 cross-study significant genes (p<0.05 using a variety of methods and datasets; **Fig. S6**; **Methods**). These diseases differ substantially between males and females in their incidence, severity, and outcomes^57,58,59^, consistent with sex-differential post- zygotic viability.

### Sex-biased study participation

So far, we have focused on forms of viability selection leading to differences in allele frequencies in our study sample because the sample represents individuals surviving to be sampled. We therefore have implicitly assumed that the study sample is representative of the true population in terms of allele frequencies. While our current approach is motivated by removing the effects of specific study recruitment biases on this assumption, it is also possible that some recruitment biases may be shared across studies. Individuals affected by diseases may be less likely or less able to participate in the study, and these diseases may differentially affect males and females. For the diseases described above (LAMA2-CMD, AML, and various cancers) as well as any diseases associated with the other cross-study significant genes, we expect that even if the disease does not cause mortality, it could affect an affected individual’s ability or willingness to share their genetic data.

Disorders not specifically affecting viability may also affect participation; for example, mental disorders may affect willingness or ability to participate. Further, any neurological or behavioral differences between males and females could affect their participation, regardless of connection to disorders or disease. Because our allele frequencies are based on only those who chose to participate in each biobank study, these biases may confound true biological selection signals. We find evidence for a gene associated with neurological and behavioral phenotypes. CNTNAP2 is associated with neuron cell organization and junction formation^63^. CNTNAP2 has also been associated with a broad range of neurological disorders, including autism spectrum disorder, schizophrenia, major depressive disorder, intellectual disability, dyslexia, and language impairment^64,65^. These disorders are known to have sexual dimorphic effects and outcomes^66,67,68,69,70^. Investigating the relationship between CNTNAP2 and sex specifically, studies have found mutations of CNTNAP2 cause sex-different effects in visual cortical brain area development^71^, dyslexia in children^72^, and disruption to social behaviors in mice^73^. Mutations to this gene may cause direct biological differences in neurobehavioral traits, leading to differences in study participation. Cultural or societal factors may further influence participation in sex-specific ways, and these factors could interact with individual genotypes at CNTNAP2 and other genes. These factors may correlate with axes of ancestry or shared environments, and these correlations may be shared across studies. Population structure confounding may in turn persist across studies^74^ through sex-differential participation. Although this neurobehavioral effect on observed allele frequencies stems from sampling methodology rather than biological viability selection, there may be genetic factors underpinning the participation bias. Disentangling these artifactual effects from viability selection will help clarify the genes that experience SDS.

## Conclusions

Sex differences in study participation as well as bioinformatic artifacts may have confounded inferences of contemporary sex-differential selection. Our meta-analysis across multiple studies can help alleviate some of these concerns. From a comparison across three genetic studies, we identify twelve genes which have concordant cross-study evidence of between-sex allele frequency divergence. Additional analyses, including of sign concordance, between-sex Hardy-Weinberg Equilibrium, haplotypic hitchhiking signals, and a literature survey offer stronger support for SDS in ANKRD36C than for the eleven other genes. Taken together, our combination of different lines of evidence provides a more robust evaluation of putative genomic targets of strong SDS in humans than previous work.

It is notable that our cross-study significant genes were not previously identified by previous studies such as Kasimatis et al. (2022)^4^, which found no SNP-level concordance across their or our biobanks. However, direct comparisons in UKB reveal significant correlation between our results and theirs for p-values and p-value ranks at the SNP and gene levels (**Fig. S7; Supplemental Methods**). The lack of exact matches at specific sites may be due to differences in data and methodology (**Supplemental Results and Discussion**). Nevertheless, the correlation between our works suggests our results are concordant with previous studies, and we offer additional criteria for identifying potential SDS candidates.

While our hypotheses about drivers of SDS remain largely speculative, we delineate the different potential modes and life stages in which SDS may act, from spermatogenesis to adulthood (**Fig. 2**) and highlight their possible connections to candidate genes. Some cross-study significant genes are associated with spermatogenesis and others with diseases with known sex differences.

To explain how spermatogenesis-related genes could be under SDS, we outlined a hypothesis for selection on individual sperm cells during fertilization. Further analysis of individual sperm to better quantify X- versus Y-sperm expression could help further elucidate this concept, but we recognize that this is a developing area of research. Recent studies such as Guo et al., 2018^75^, have analyzed the testes of adult males using single-cell RNAsequencing (scRNA-seq) to better understand transcription in different cell types, including sperm. However, the authors note that sperm and sperm precursors are known to have poor RNA content and thus are less reliable for analysis^75^. Future work building upon higher quality scRNA-seq sperm data could help further development of this hypothesis. We also acknowledge that this hypothesis is backed by associative evidence of genes associated with spermatogenesis and transcription patterns, and further functional analysis of specific genes will clarify their roles in contributing to SDS.

Several study-significant genes were consistent with the hypothesis of sex-differential post-zygotic viability, the sole or major driver of SDS highlighted by previous work^1,2,4,5^.

Additionally, although testing for cross-study signals may help mitigate effects of dataset-specific biases, some recruitment biases may persist across studies. We find cross-study evidence for a gene that affects neurological disorders. While these disorders may impact viability, it is likely that these and other health conditions affect study participation to a sex-differential extent.

In total, this work helps delineate drivers of sex differences in allele frequency and finds a small number of putative targets with cross-study evidence consistent with these hypothesized drivers. Methodologically, we suggest improvements to existing standards of testing, interpretating and characterizing genetic signals of SDS. Nevertheless, allele frequency differences in retrospective studies are inherently difficult to interpret, and the hypotheses we put forth here—or future ones based on similar approaches and data—should be treated solely as preliminary hypotheses to guide further research.

## Methods

### Whole genome sequence data from three genetic studies

We used data from the UK Biobank (UKB), All of Us (AoU), and gnomAD datasets. To obtain a list of samples for UKB, we first filtered individuals by their alignment with the White British cohort by principal component analysis. Similarly, for AoU, we again filtered individuals by ancestry matching with the White British cohort in UKB using a random forest classifier^74^. In gnomAD, individuals are pre-separated into genetic ancestry groups by principal component analysis. We used individuals in the Non-Finnish European (NFE) ancestry group as the closest ancestry match to UKB. For all studies, we filtered and identified male and female samples by XY or XX chromosomal sex, respectively. After ancestry matching and sex chromosome filtering, we were left with 186,936 males and 220,062 females in UKB, 79,983 males and 124,205 females in AoU, and 14,343 males and 19,686 females in gnomAD.

To obtain allele counts in males and females, we used whole genome sequences in each dataset. In gnomAD, we used VCFtools to obtain allele counts and sample numbers from the whole-genome sequence summary statistic VCF files from gnomAD v3 available for download, accessed in June 2024. For allele counts in UKB, we used data from the 500k Whole Genome Sequence (WGS) release (accessed January 2025). We obtained allele and genotype counts for males and females separately from 20kbp blocks using PLINK^76^, then concatenated these blocks for each chromosome. For allele counts for AoU, we used data from the 250k WGS release (accessed February 2025). Here, we focused on obtaining allele counts from individual genes, specifically a set of genes identified as having significant allele frequency divergence in both UKB and gnomAD, and a set of random non-significant genes to act as a null expectation (see Methods: Chi-square test for allele frequency divergence). We obtained allele and genotype counts for this set of genes using the plink2 command from PLINK2 v. 2.0^76^.

We filtered SNPs to only include biallelic autosomal SNPs. We also performed a BLAST^77^ mapping between 150bp sequences surrounding ascertained SNPs and the X and Y chromosomes. SNPs with over 90% identity with X and Y chromosome sequences were excluded from the study. To further mitigate potential mismapping to sex chromosomes, we filtered out sites which lay within segmental duplicates with X/Y paralogs^24,78^. Further analysis of sequence identity for segmental duplicates specifically within cross-study significant genes and paralogs ANKRD36 and ANKRD36C was conducted using BLAST. Finally, we filtered sites to only those within gene bodies (as annotated in Gencode v21) as well as 1000bp upstream to include promoters.

### Chi-square test for cross-study allele frequency divergence

To test for significant between-sex allele frequency divergence, we performed a chi-square test. The 2x2 contingency table was constructed with male and female major and minor allele frequencies. We performed this chi-square test on all SNPs in the genome in UKB and gnomAD separately. To measure the gene-level significance, we took the single lowest p-value site within a gene as the p-value for the gene as a whole, and we generated a distribution of gene p-values. Because we were most interested in the most significantly diverged genes in each population, we used an outlier approach to identify cross-study significant genes. For a given genetic study, we identified “significant genes” as those whose lowest p-value SNP is within the bottom 1% of genes in that study. Genes which were in the bottom 1% in both datasets were considered “overlapping significant genes”.

To identify significant genes in AoU, we specifically focused on the overlapping significant genes from UKB and gnomAD. We performed the chi-square test on all SNPs in the overlapping significant genes, as well as in 200 randomly selected genes to form the null distribution. Overlapping significant genes whose p-values were below the 1% threshold for the 200 randomly selected null gene p-values were considered “cross-study significant genes”, where their p-values were outliers in all three datasets. We identified 12 such genes ( **Table 1**).

### MAGMA analysis for gene-wide p-values

To account for gene length, we also computed p-values for genes using MAGMA. To do this, we performed a MAGMA analysis following methods and the magma command from software presented by De Leeuw et al., 2015^18^. MAGMA generates a test statistic which aggregates all SNPs within a gene and can be performed using per-SNP p-value summary statistics. For generating a SNP correlation matrix, we used the genotype reference panel from 1000 Genomes phase 3, European ancestry group^79^. For our analysis, we used the chi-square test p-values calculated for all SNPs in a gene as above (see **Methods: Chi-square test for cross-study allele frequency divergence**). In short, the MAGMA method performs a weighted sum of the -log_10_(p-values) to generate a MAGMA test statistic *T*, where the weights are based on SNP correlations. This *T* follows a gamma distribution, with shape parameter *α* = *E*(*T*)/*Var*(*T*) and scale parameter θ = *Var*(*T*)/*E*(*T*), where 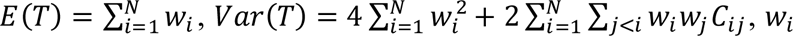 are the weights, *C_ij_* is the covariance between SNP i and SNP j determined from the SNP correlation matrix, and *N* is the number of SNPs in the gene. A gene-wide p-value is then generated by finding the probability of *T*∼Γ(*α*, *θ*).

### Male-Female allele frequency difference pairwise sign concordance

To calculate concordance of sign, we investigated the between-sex difference in allele frequency at individual SNPs. For each dataset, for each cross-study significant gene, a single SNP was the site with the lowest p-value; in the case that multiple SNPs had equally low p-values, we used the site with the greatest difference in allele frequency. Because each dataset possibly had a different SNP as the lowest for a given gene, we looked at three different SNPs per gene, termed the “UKB site”, “gnomAD site”, and “AoU site” respectively. We first polarized alleles by which one had the higher frequency in males in UKB. We then calculated the male-minus-female difference in allele frequency in all three datasets separately and took the sign from that difference; a positive sign meant the allele was more frequent in males, and negative meant more frequent in females. The same sign between two datasets was recorded as a concordant sign. We compared the sign of the UKB site, gnomAD site, and AoU site separately, and recorded the concordance in **Table S1**.

### Linked site cross-study sign concordance

To account for LD causing neighboring sites to diverge in frequency and potentially causing us to under-capture sign concordance between studies, we also performed a comparison of the SNP in strongest LD with the lead SNP in each study. To find the strongest LD SNP, we calculated LD of the lead SNPs with all other SNPs in the gene body using the UKB reference panel for UKB, and the AoU reference panel for AoU; LD in gnomAD was also calculated using the AoU reference panel because individual genotype data is not available for gnomAD v3. We then summed these LD values for all three lead SNPs. The SNP with the highest sum LD was considered the strongest LD SNP. We then compared the sign of male-female allele frequency divergence at this SNP in each study, again polarizing by allele with the higher frequency in males in UKB.

### Test for deviation from Hardy-Weinberg Equilibrium

HWE tests require genotype count data and sex-specific genotype frequencies were not available through gnomAD v3. We therefore tested for deviations from HWE only in UKB and AoU. To test for deviation from Hardy-Weinberg Equilibrium, we used a formula from Wigginton, Cutler, and Abecasis, 2005^26^. The formula for the probability of exactly the observed number of heterozygotes given a sample size *N* and number of *A* alleles *n_A_* is as follows (copied from their paper, equation 1):

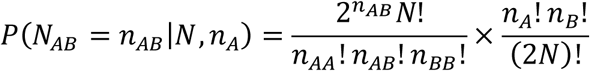

To calculate the p-value for observing this or a more extreme number of heterozygotes under HWE, we then compared the probability of observed heterozygotes with the probability of all possible numbers of heterozygotes and sum the probabilities less than or equal to the observed heterozygote probability.

Because our sample sizes were very large for some sites (on the order of many tens or hundreds of thousands), calculating an exact factorial became impractical. Therefore, we used Ramanujan’s factorial approximation, where 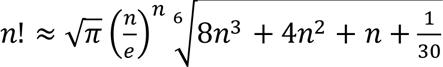 and by extension 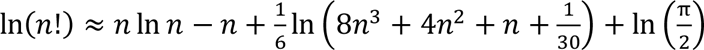 So 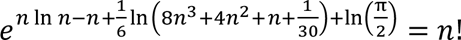

### Comparison with genotype informative markers in sperm

Genes were identified as being expressed in sperm if they were captured in the data and results from Bhutani et al., 2021. They were identified as not being shared across bridges if they were identified as Genotype Informative Markers (GIMs) in the same study. The enrichment p-value for genes expressed in sperm was calculated using a Fisher’s exact test, using the formula

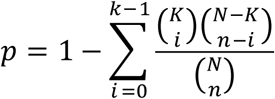

Where *N* is the total number of protein-coding genes in our annotation, *K* is the total number of genes found expressed in sperm, *n* is the total number of cross-study significant genes we identified, and *k* is the total number of cross-study significant genes found to be expressed in sperm.

### Disease enrichment in cross-study significant genes

To test for enrichment of sex-differential diseases among the cross-study significant genes, we performed a gene-set enrichment analysis using WebGestalt^80^. This web-based analysis tool sources from multiple association databases, including databases specifically for disease-gene associations. We compiled results from two databases: Online Mendelian Inheritance in Man (OMIM)^81^ and DISGENET^82^.

We performed enrichment analysis in two ways. First, we compared the diseases associated with the set of 12 cross-study significant genes against all protein-coding genes as a background set. Second, we used a single list of genes ranked by chi-sq p-value, looking for enrichment of diseases associated with genes near the top of the list compared to the bottom.

## Code Availability

Scripts used in this study are available at https://github.com/harpak-lab/SDS_Drivers. Raw data from each biobank is available either for free download online (gnomAD: https://gnomad.broadinstitute.org/data) or through the respective online workspaces (UKB RAP and AoU Workbench 2.0). Analyzed data for downstream analysis is available at the Zenodo repository https://doi.org/10.5281/zenodo.21478252

## Acknowledgements

We thank Mark Kirkpatrick for many helpful comments throughout the analysis and writing process. We thank Ipsita Agarwal and members of the Harpak lab for helpful conversations. We thank the Texas Advanced Computing Center for providing computational resources that have contributed to the research results published in this paper. This project was conducted using the UK Biobank and UK Biobank Research Analysis Platform under application number 61666. We thank the National Institutes of Health’s All of Us Research Program for making available the participant data examined in this study. We gratefully acknowledge the gnomAD, UK Biobank, All of Us participants for their contributions, without whom this research would not have been possible. We thank the Institutional Review Board for oversight and commitment to ethical research. This research was funded by NIH grant GM151108 and a Pew Scholarship to A.H.

