## Supplemental Information for "Drivers of systemic male-female allele frequency divergence in humans"

Fig. S1: Distributions of p-values in gnomAD, UKB, and AoU

Fig. S2: Allele frequency differences in cross-study significant genes that were not significant by *MAGMA*

Fig. S3: Cross-study sign concordance of lead SNPs

Fig. S4: Gene-level p-value tends to decrease as gene length increases

Table S1: Gene-wide aggregated p-values calculated using *MAGMA*

Fig. S5: Comparison of deviations from Hardy-Weinberg equilibrium for most significant SNP in cross-study significant genes

Table S2: Genes putatively under SDS which are related to spermatogenesis and fertility are also among those that are not shared across cytoplasmic bridges

Fig. S6: Enrichment of diseases among 12 cross-study significant genes

### Supplemental Methods

<u>Comparison with Kasimatis et al.</u>

We compared our results with Kasimatis et al. (2020). Notably, the sites identified by Kasimatis et al. had no overlap with our own results, and we sought to understand how much our results differ. We used the UK Biobank data in both studies as a baseline comparison point. We used Kasimatis et al.'s genetic sex GWAS p-values for each site. SNP-level correlations between our results were performed by first filtering only to include SNPs which were ascertained by both our studies. Then we compared the Kasimatis et al p-values with our chi-square p-values using a Pearson correlation test. We also ranked each SNP by p-value and re-calculated the correlation using these ranks.

To compare gene-level results, we used the Ensembl gene annotations to find all Kasimatis et al. SNPs within the gene body or 1000bp upstream (as in the main text **Methods:** "Whole genome sequence data from three genetic studies"). We then took the lowest p-value of SNPs within a given gene as the gene-wide p-value. As with the SNP level analysis above, we performed a Pearson's correlation test on these gene-level p-values and the ranks of the p-values.

### Supplemental Results and Discussion

We find there is significant agreement between our study and Kasimatis et al., particularly at the gene level which was the primary unit of our analysis. Differences in methodology, and specifically in sequencing and SNP ascertainment, biobank differences, and differences in sample filtering by ancestry could account for the lack of perfect agreement in our results.

We compared our UKB results to those of Kasimatis et al. (2020) by calculating correlation in four metrics: individual SNP p-values (**Fig. S7a**) and p-value ranks (**Fig. S7b**); and in the p-values and ranks at the gene level (**Fig. S7c-d**). All of these correlations were significant and positive, with the significance and strength of correlation when looking at ranks compared to p-values, and when looking at genes compared to SNPs.

We do recognize the correlation is not exactly 1:1, and the specific SNPs identified as significant do not overlap. This discordance may be due to three factors: 1) Kasimatis et al. used array SNPs whereas we used whole-genome sequence (WGS) data, leading to potentially different SNPs being ascertained; 2) our UKB and AoU sample sizes are larger than Kasimatis et al.'s UKB and BioVU (although the gnomAD sample sizes are smaller); and 3) Kasimatis et al. used somewhat different sample and SNP filtering procedures.

Overall, differences in data and methodology exist between our and the Kasimatis et al. study, but despite this we see significant concordance between our results. We are especially encouraged by the concordance we see at the gene level.

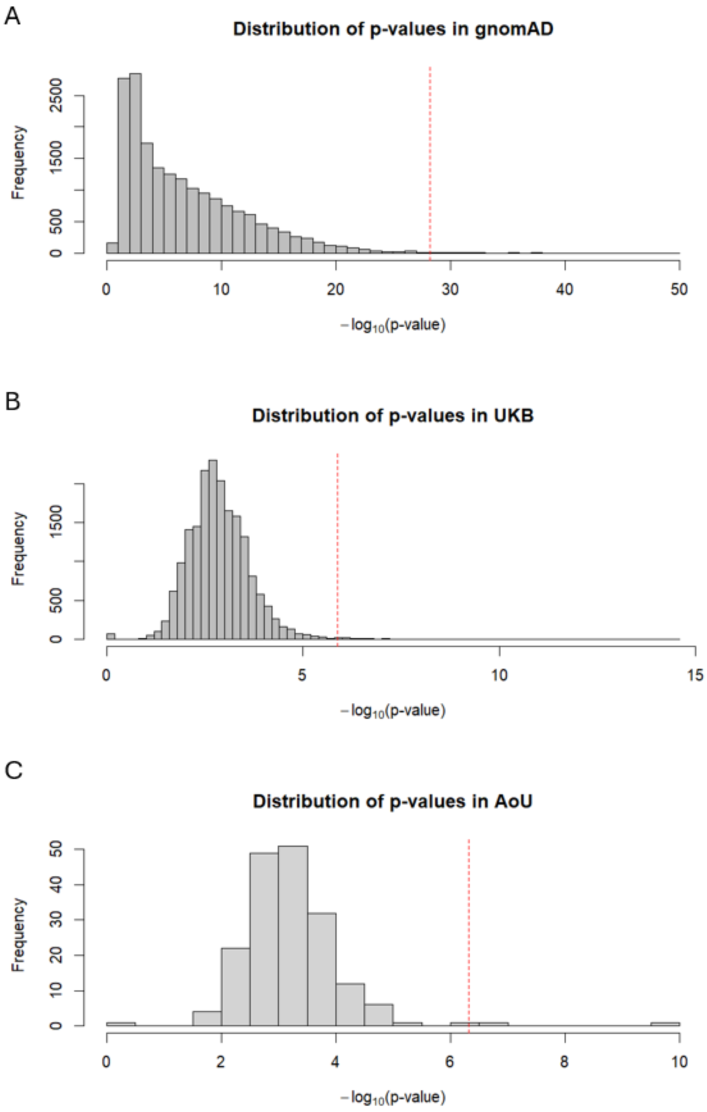

**Supplemental Figure 1: Distributions of p-values in gnomAD, UKB, and AoU.** The dotted lines show the 99% cutoff used for determining cross-study significant genes. The AoU distribution is based on a matched null compared to a set of cross-study significant genes across gnomAD and UKB.

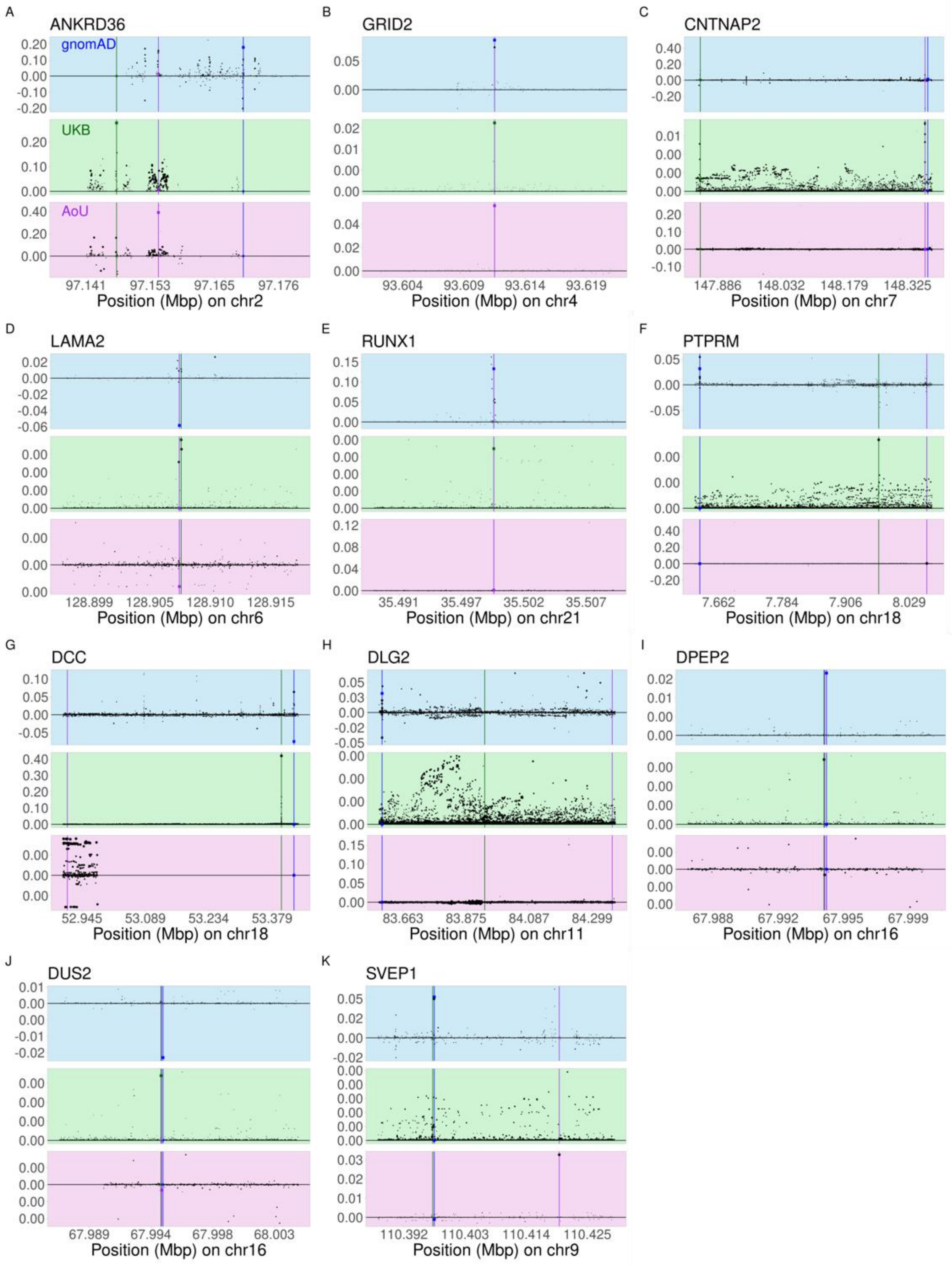
A
ANKRD36
gnomAD
UKB
AoU
97.141 97.153 97.165 97.176
Position (Mbp) on chr2
B
GRID2
93.604 93.609 93.614 93.619
Position (Mbp) on chr4
C
CNTNAP2
147.886 148.032 148.179 148.325
Position (Mbp) on chr7
D
LAMA2
128.899 128.905 128.910 128.915
Position (Mbp) on chr6
E
RUNX1
35.491 35.497 35.502 35.507
Position (Mbp) on chr21
F
PTPRM
7.662 7.784 7.906 8.029
Position (Mbp) on chr18
G
DCC
52.945 53.089 53.234 53.379
Position (Mbp) on chr18
H
DLG2
83.663 83.875 84.087 84.299
Position (Mbp) on chr11
I
DPEP2
67.988 67.992 67.995 67.999
Position (Mbp) on chr16
J
DUS2
67.989 67.994 67.998 68.003
Position (Mbp) on chr16
K
SVEP1
110.392 110.403 110.414 110.425
Position (Mbp) on chr9

**Supplemental Figure 2: Allele frequency differences in cross-study significant genes that were not significant by *MAGMA*.** We identified eleven genes that do not show *MAGMA* significance but do show cross-study significance by Fisher's exact test. As in Fig. 1, each point is a single SNP, with its size reflecting Chi-squared test p-value. The y-axis shows male-minus-female allele frequency, polarized by the allele with higher frequency in males in UKB. Plots are colored by the dataset they show (blue for gnomAD, green for UKB, purple for AoU). Colored square points represent the lowest p-value site in each dataset.

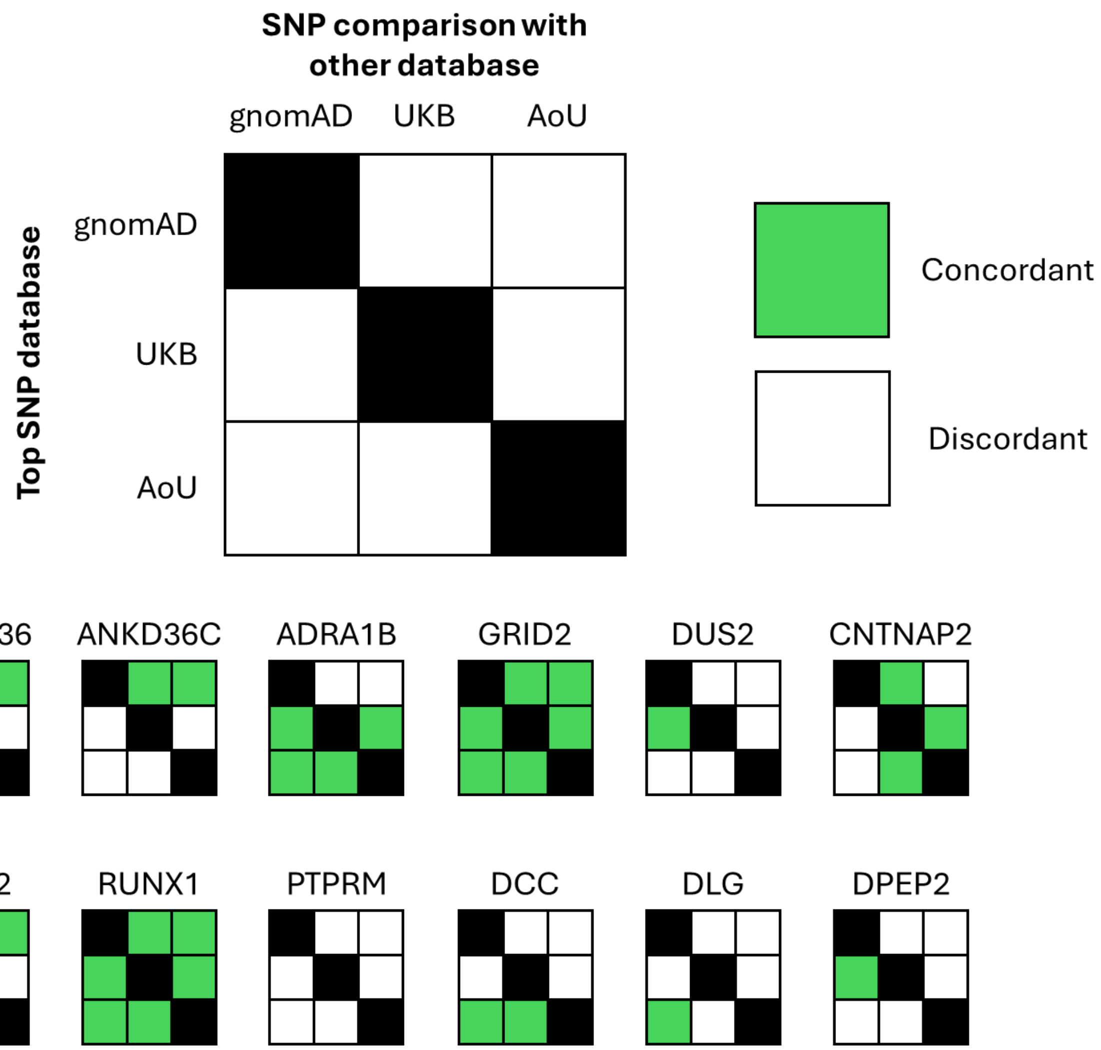

**Supplemental Figure 3: Cross-study sign concordance of lead SNPs.** We report the pairwise sign concordance for the most significant SNP in each study. The study from which a given lead SNP was obtained is in each row, and the study with which the sign is compared is in each column. Cells are colored green if the sign is concordant, and white if the sign is discordant.

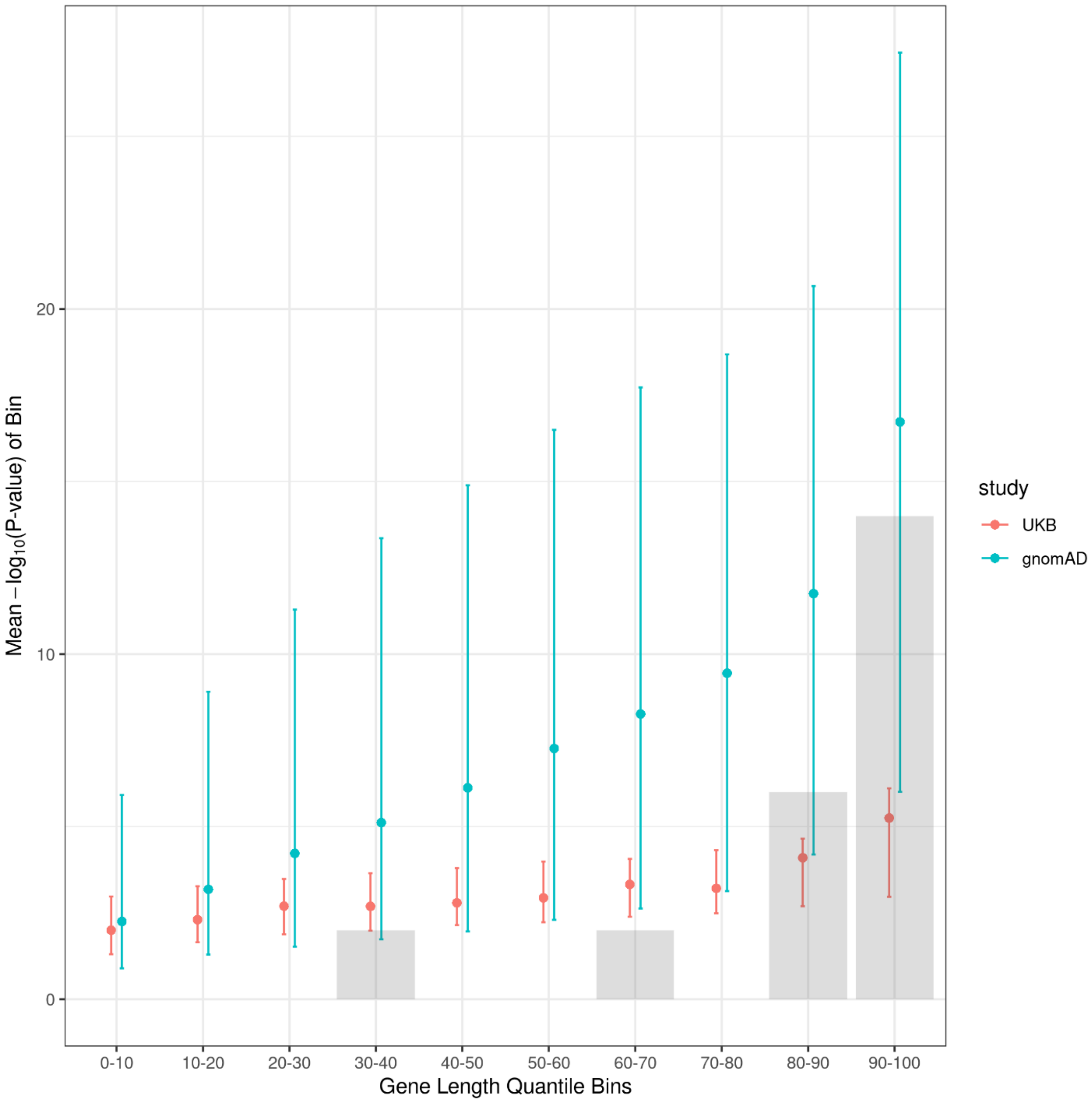

**Supplemental Figure 4: Gene-level p-value tends to decrease as gene length increases.** Cross-study significant genes are enriched for long gene lengths. The x-axis is gene size binned into ten equally sized bins by quantile. Teal and red points represent the means in gnomAD and UKB respectively, while the blue and red lines represent the 5th to 95th confidence intervals for those datasets. The grey boxes represent relative abundances of the cross-study significant genes in each quantile bin.

| GENE | P-value (gnomAD) | P-value (UKB) | P-value (AoU) |
|---|---|---|---|
| ANKRD36C | 1.33e-03 | 2.03e-06 | 1.09e-07 |
| ANKRD36 | 0.12 | 4.83e-03 | 4.40e-03 |
| GRID2 | 0.95 | 0.71 | 3.257e-03 |
| DCC | 0.71 | 0.12 | 3.10e-02 |
| DUS2 | 0.66 | 0.33 | 0.06 |
| SVEP1 | 0.26 | 0.41 | 0.90 |
| DLG2 | 0.34 | 0.64 | 0.55 |
| PTPRM | 0.30 | 0.47 | 0.92 |
| CNTNAP2 | 0.76 | 0.76 | 0.56 |
| RUNX1 | 1.00 | 0.75 | 0.77 |
| LAMA2 | 0.97 | 0.85 | 0.80 |
| DPEP2 | N/A | N/A | 0.06 |

**Supplemental Table 1: Gene-wide aggregated p-values calculated using *MAGMA*.** P-values represent the probability of a summary statistic calculated by summing p-values calculated for all SNPs within a gene, weighted by their correlation with each other. Cells are colored by their significance, with darkest being most significant ($p < 0.001$) and lightest being least ($p < 0.05$).

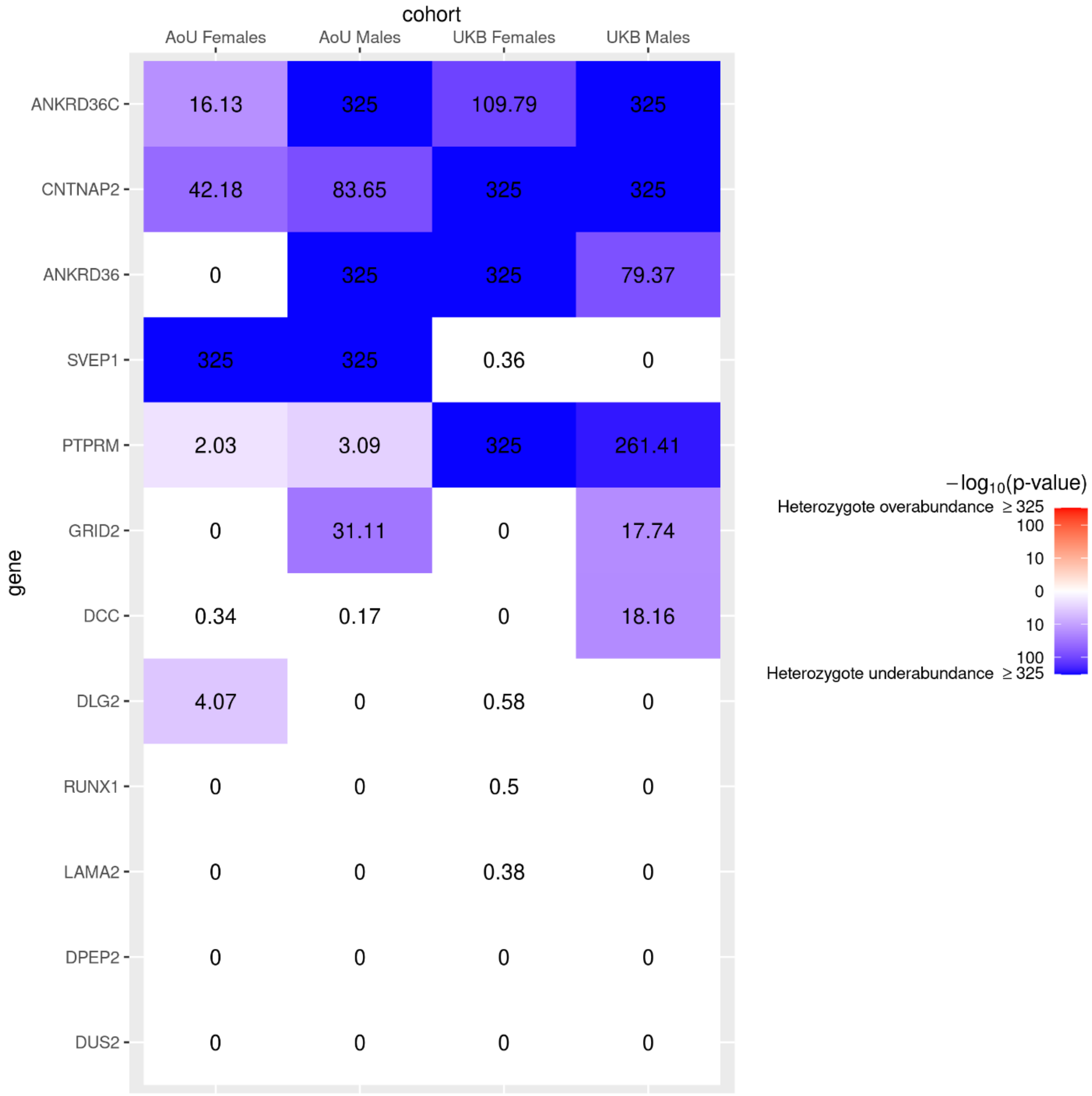

**Supplemental Figure 5: Comparison of deviations from Hardy-Weinberg equilibrium for most significant SNP in cross-study significant genes.** Color intensity represents p-value for deviation from Hardy-Weinberg Equilibrium (HWE) by Fisher's Exact Test, with darker colors corresponding to more significant deviations from the null of concordance with HWE (i.e., no selection).

| Gene Name | Shared across bridges |
|---|---|
| GRID2 | Not shared across bridges |
| ANKRD36C | Not shared across bridges |
| DCC | Not shared across bridges |
| ANKRD36 | Shared across bridges |
| DUS2 | No data |
| PTPRM | Non-significant results |

**Supplemental Table 2: Genes putatively under SDS which are related to spermatogenesis and fertility are also among those that are not shared across cytoplasmic bridges.** Based on results from Bhutani et al. (2021), several of the cross-study significant genes associated with spermatogenesis and fertility are also not shared across cytoplasmic bridges during spermatogenesis, indicating that these genes could have direct phenotypic effects of individual sperm cell genotype on individual sperm cell phenotype.

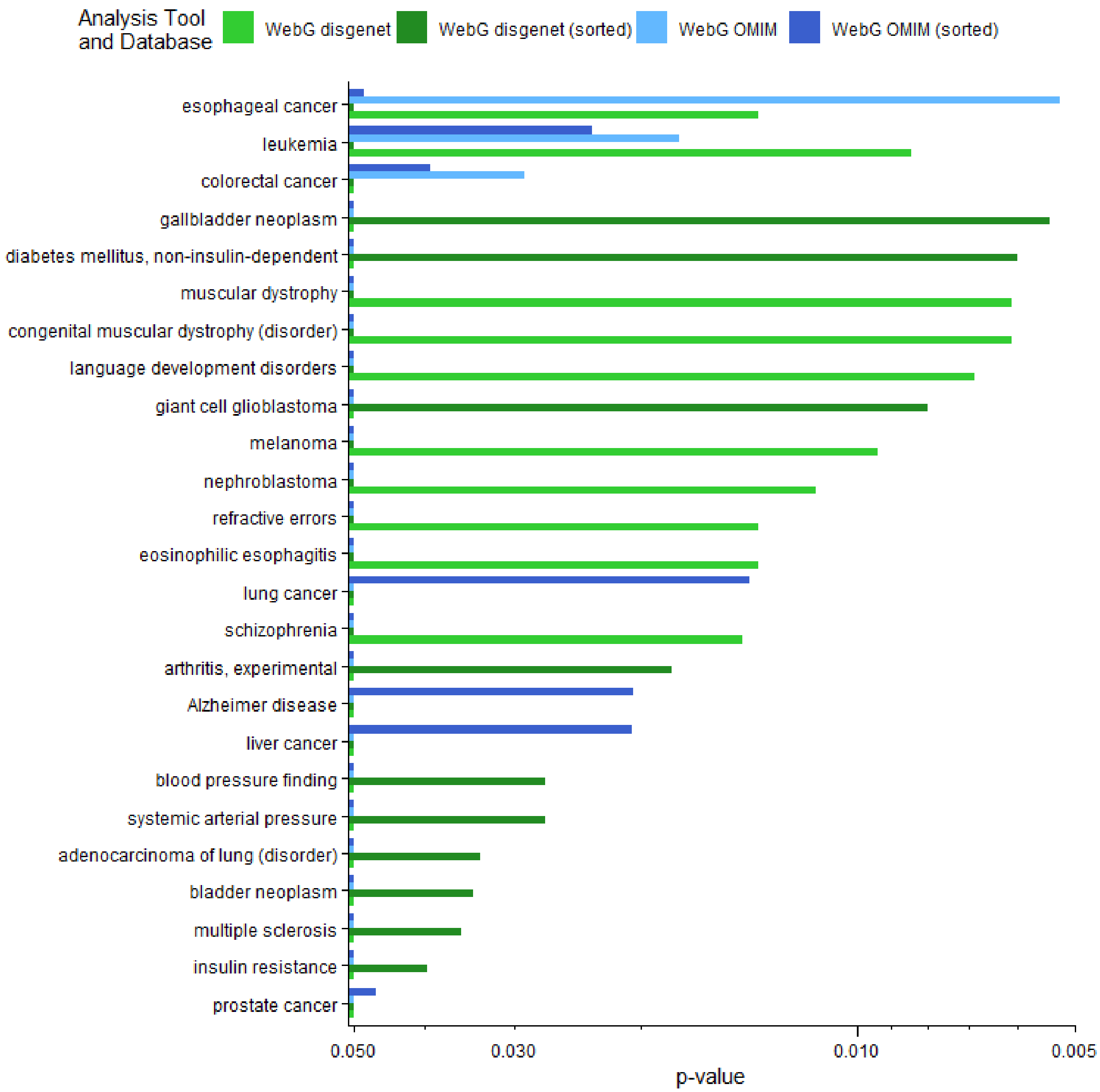

**Supplemental Figure 6: Enrichment of diseases among 12 cross-study significant genes.** Bars are colored by methodology used for enrichment analysis. Only significantly enriched diseases with $p < 0.05$ are shown.

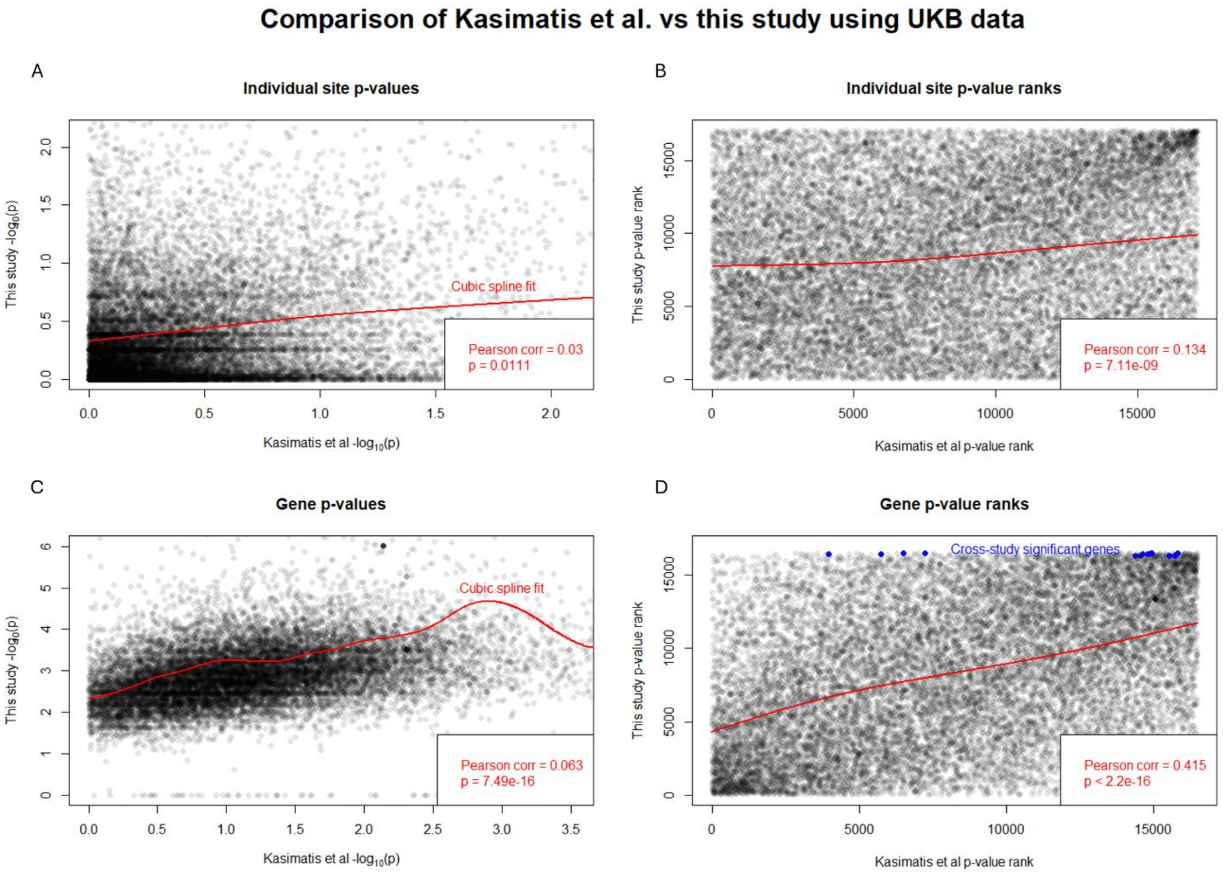

**Supplemental Figure 7: Comparison of Kasimatis et al. p-value results with present study.** We compared our results with those obtained by Kasimatis et al. (Genetics, 2020) for UKB. Shown are comparisons between individual site p-values (A), individual site p-value ranks (B), gene-level p-values (C), and gene-level p-value ranks (D). In all panels, the red line shows a cubic spline fit to the data. In panel D, the blue points show the comparisons of the cross-study significant genes identified in the present study.
